## Supplementary Information for "Interacting with volatile environments stabilizes hidden-state inference and its brain signatures"

**Contents:**

Supplementary figures 1-11 (pages 2-12)
Computational model specifications (pages 13-20)
Supplementary references (page 21)

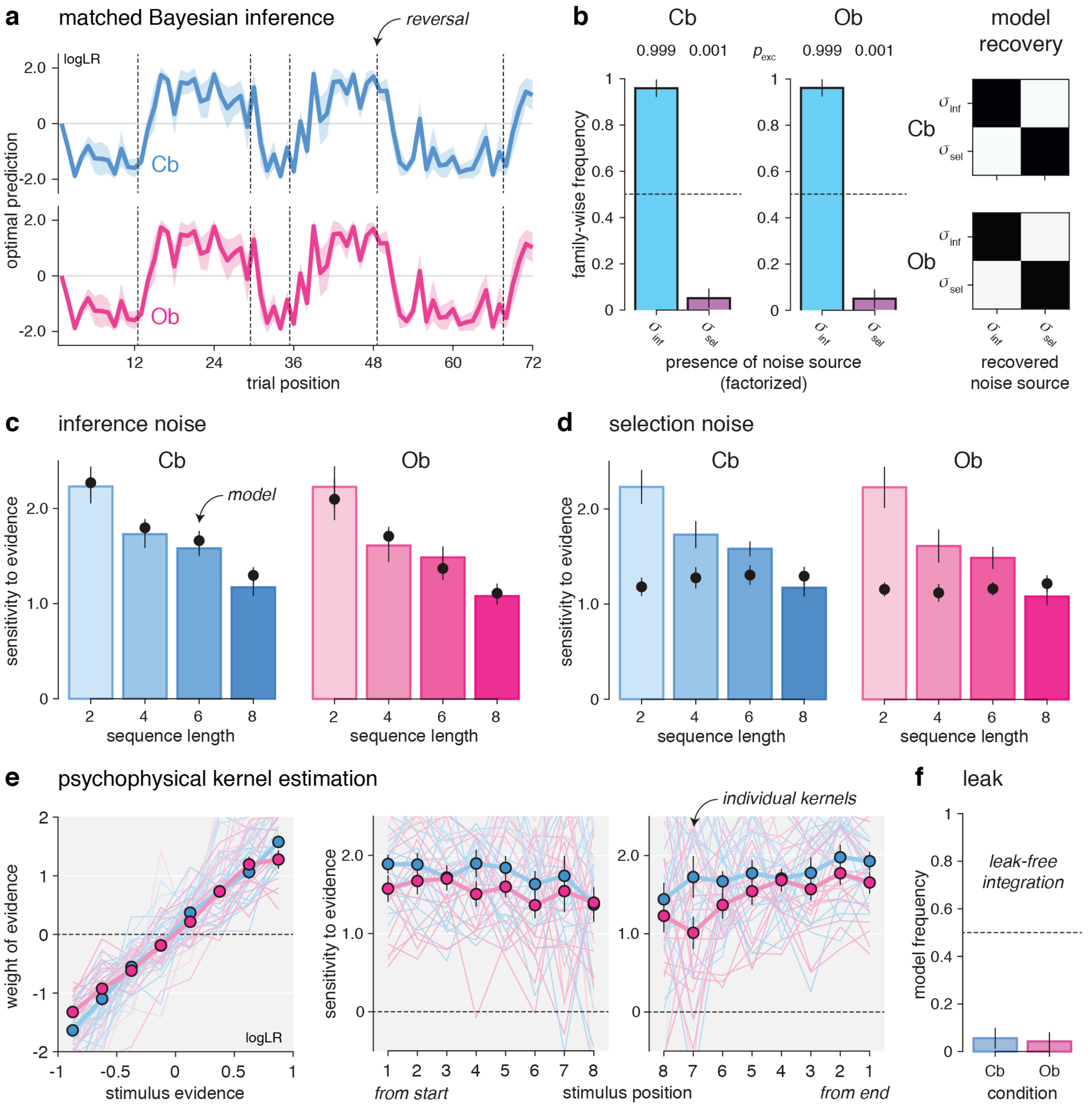


**Supplementary Fig. 1 – Matched suboptimal Bayesian inference across conditions. a,** Matched trajectories of Bayes-optimal predictions (beliefs at the onset of each trial, expressed as log-odds ratio) for two matching blocks in the cue-based and outcome-based conditions. The trajectories of beliefs shown (obtained using $\sigma_{\mathrm{inf}}$ = 0.5 and $h$ = 0.125) were matched by using the same pre-generated stimulus tilts for the two blocks. **b**, Bayesian model selection regarding the sources of behavioral variability in the cue-based and outcome-based conditions. Left: the left bar shows the estimated frequency of inference noise ($\sigma_{\mathrm{inf}} > 0$), whereas the right bar shows the estimated frequency of selection noise ($\sigma_{\mathrm{sel}} > 0$). Exceedance probabilities $p_{\mathrm{exc}}$ are reported above each bar. Participants feature significant inference noise but no selection noise. Right: model recovery results regarding the sources of behavioral variability in the cue-based and outcome-based conditions. Confusion matrices show the estimated frequency (from white = 0 to black = 1) of each noise source (columns) for simulations of each noise source (rows). The two noise sources can be accurately identified through Bayesian model selection. **c,** Validation of inference noise as main source of behavioral variability. Simulations of inference noise predict that the sensitivity of participants' decisions to evidence should decrease as a function of the number of inference steps in a given trial (black dots). Participant estimates (bars) match model simulations in both conditions. **d**, Falsification of selection noise as main source of behavioral variability. Simulations of selection noise predict that the sensitivity of participants' decisions to evidence should be independent of the number of inference steps (black dots). Participant estimates (bars) deviate from model simulations in both conditions. **e**, Psychophysical kernel estimation. Left: unbiased weighting of evidence. Estimated weight of evidence (y-axis) as a function of objective stimulus evidence (x-axis) binned into eight equally-spaced bins. Dots and error bars indicate group-level means ± s.e.m. Right: leak-free accumulation of evidence. Estimated sensitivity to stimulus evidence (y-axis) as a function of stimulus position from start (left) and end (right) of a sequence (x-axis). **f**, Bayesian model selection regarding the presence of an integration leak across stimuli of the same sequence. Participants integrate evidence without detectable leak in both conditions.

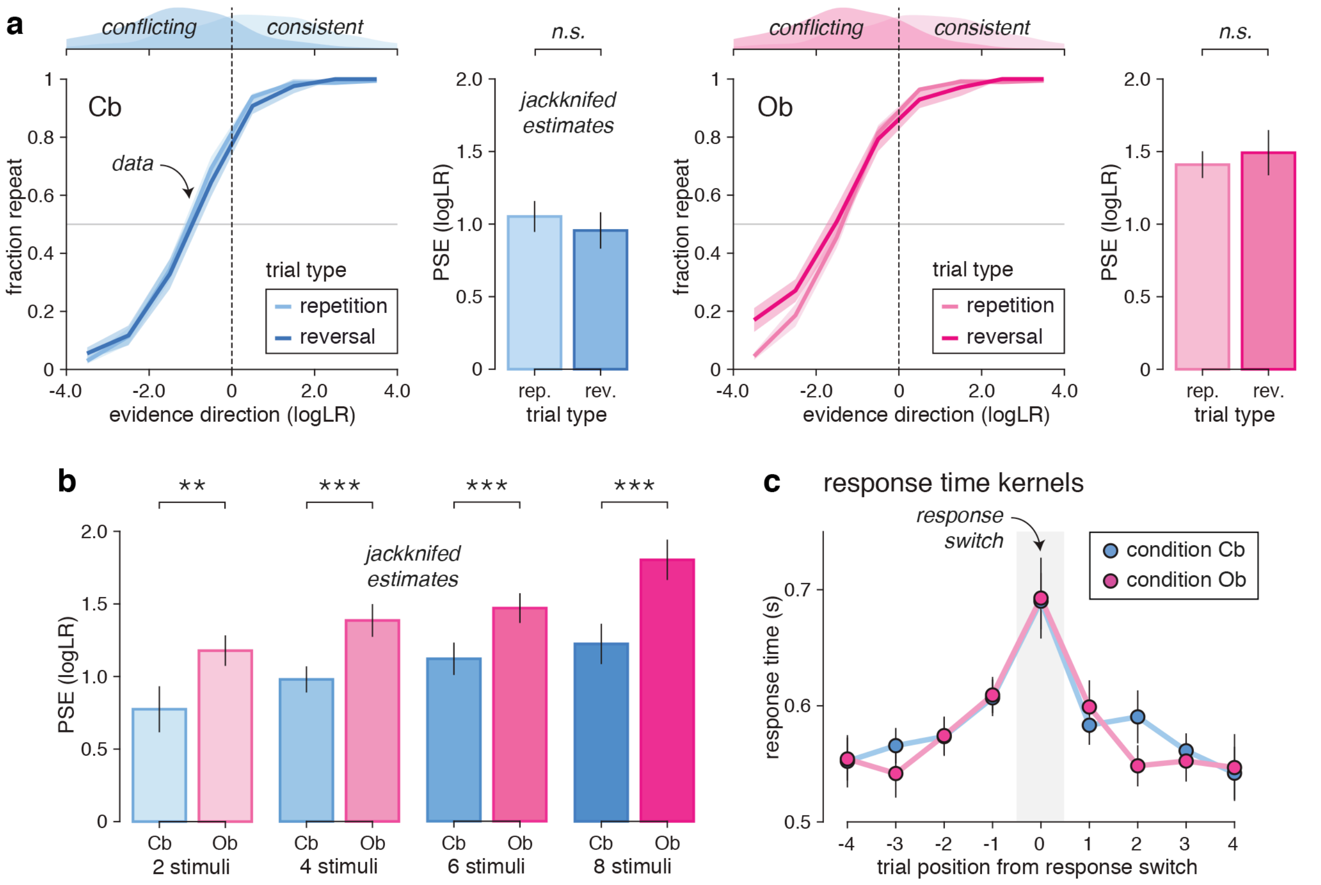


**Supplementary Fig. 2 – Robustness of psychometric effects to trial-to-trial variability in task parameters. a**, Response repetition curves in the cue-based (left) and outcome-based (right) conditions, split between trials where the hidden state $s$ either repeats itself (lighter colors) or reverses (darker colors). Lines and shaded error bars indicate jackknifed group-level means ± s.e.m. Although trials where the hidden state reverses are associated with more conflicting evidence than trials where the hidden state repeats itself (top insets), these two classes of trials are associated with identical PSE estimates (right insets). Bars and error bars indicate jackknifed group-level means ± s.e.m. **b**, Best-fitting jackknifed estimates of PSE as a function of the number of stimuli in the current sequence. Bars and error bars indicate jackknifed group-level means ± s.e.m. The increased PSE in the outcome-based condition is robust to trial-to-trial variability in sequence length. **c**, Matched response time kernels surrounding response switches. Dots and error bars indicate group-level means ± participant-level s.e.m. Response switches (and trials preceding/following switches) are associated with increased response times in both conditions. Response switches are associated with identical response times in the two conditions, in disagreement with a response-level account of the difference between conditions. Two stars indicate a significant effect at $p < 0.01$, three stars at $p < 0.001$, *n.s.* a non-significant effect.

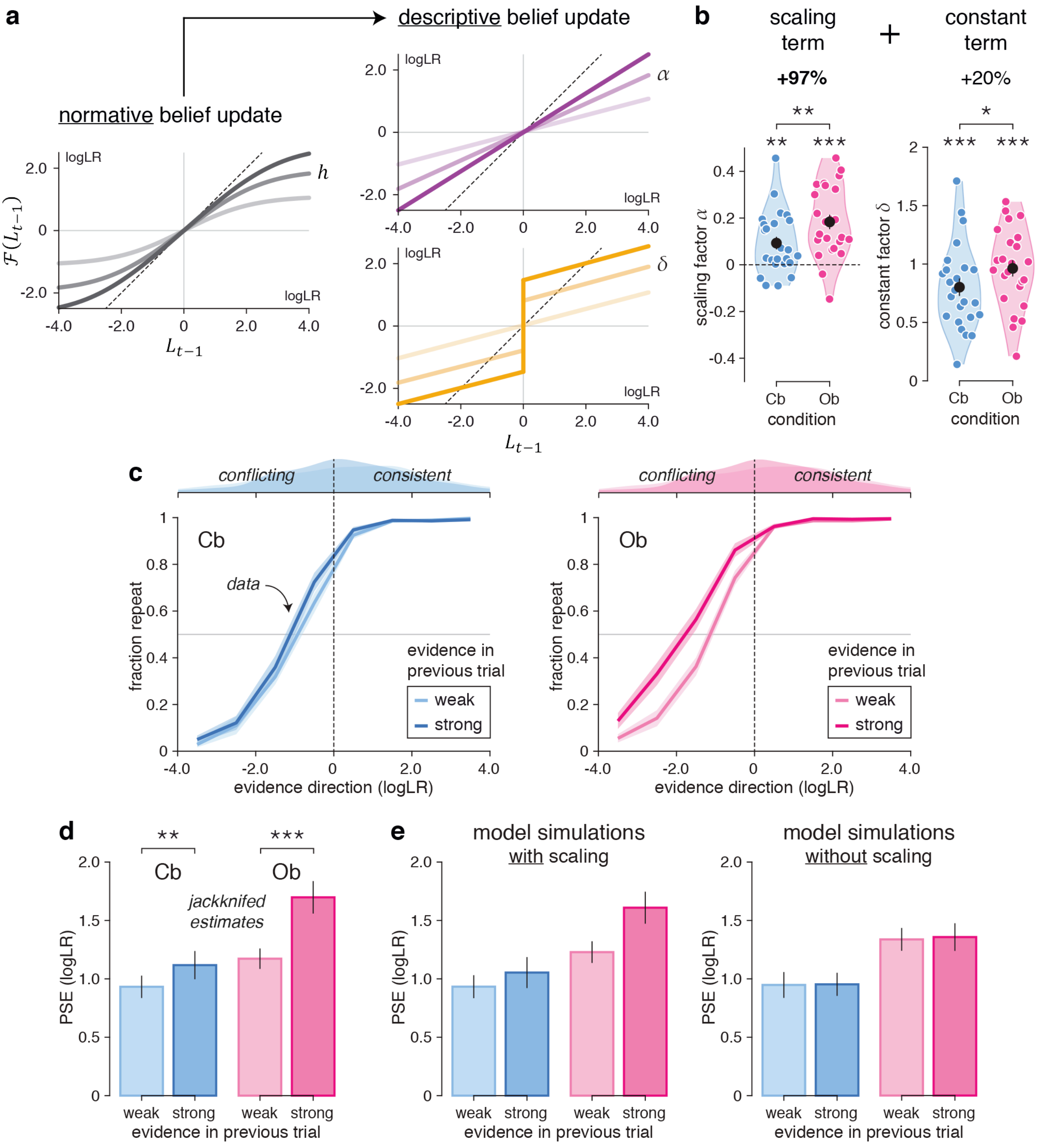


**Supplementary Fig. 3 – Descriptive modeling of hidden-state inference.** **a**, Descriptive update of the posterior belief $L_{t-1}$, controlled by two free parameters: a multiplicative scaling factor $\alpha$ (top), and an additive constant factor $\delta$ (bottom). This descriptive update decomposes the normative update (left) into a scaling (gain) term, and a constant (bias) term. **b**, Best-fitting parameter values for the scaling factor (left) and the constant factor (right). Black dots and error bars indicate group-level means ± s.e.m., whereas colored dots indicate participant-level estimates. The scaling factor increases by 97% in the outcome-based condition, whereas the constant factor increases only by 20%. **c**, Response repetition curves in the cue-based (left) and outcome-based (right) conditions, split between trials where the evidence provided by the previous sequence in favor of the previous response is either weak (smaller than its median value, lighter colors) or strong (larger than its median value, darker colors). Lines and shaded error bars indicate jackknifed group-level means ± s.e.m. Strong evidence in the previous trial in favor of the previous response shifts psychometric curves leftwards. **d**, Best-fitting jackknifed estimates of PSE as a function of evidence in the previous trial. Bars and error bars indicate jackknifed group-level means ± s.e.m. Strong evidence in the previous trials increases the PSE significantly more in the outcome-based condition. **e**, Predicted jackknifed estimates of PSE for simulations of the best-fitting model either with a scaling term (left) or without a scaling term (right). Model simulations with a scaling term can account for the observed interaction between condition and evidence in the previous trial, whereas model simulations without a scaling term fail to account for this effect. One star indicates a significant effect at $p < 0.05$, two stars at $p < 0.01$, three stars at $p < 0.001$.

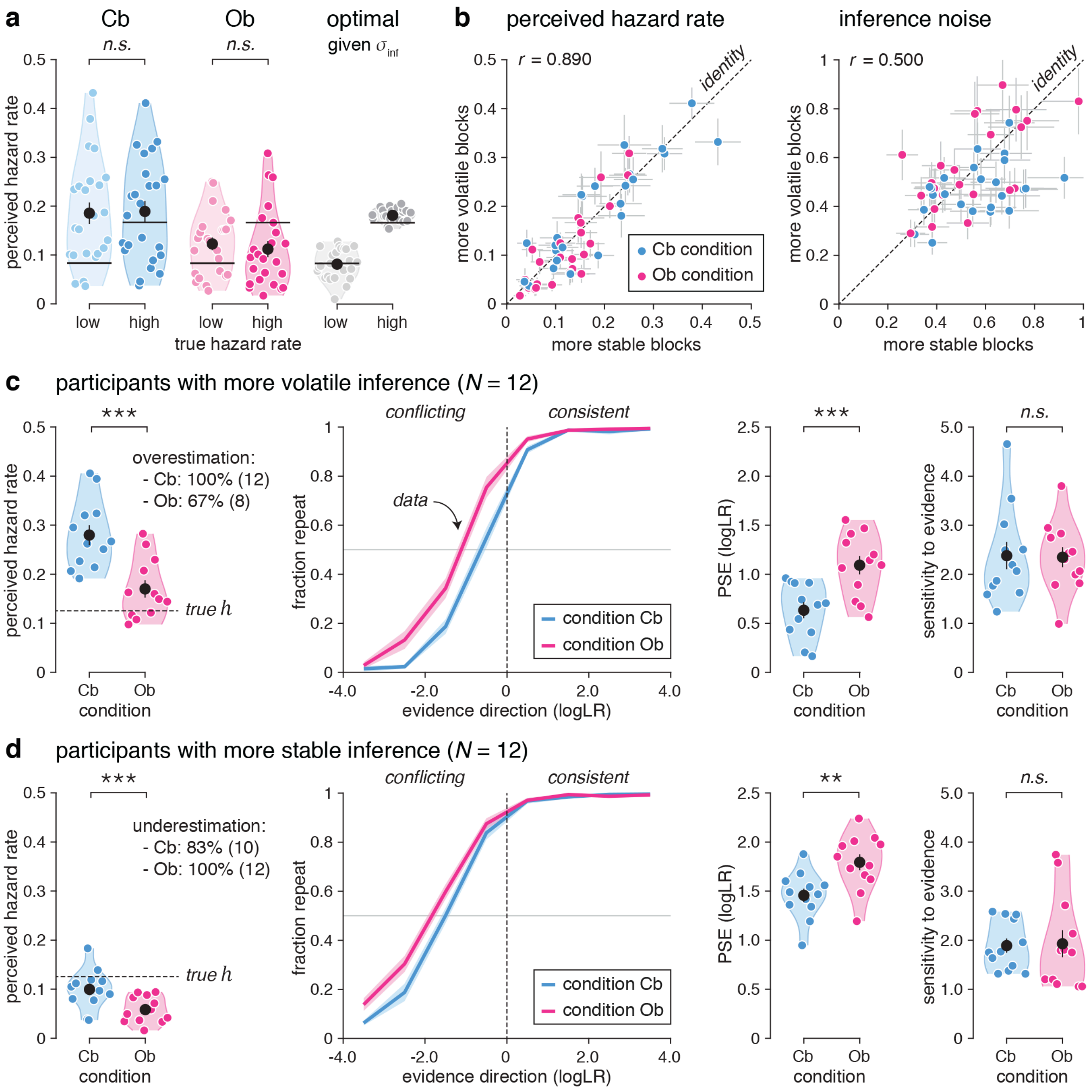


**Supplementary Fig. 4 – Robustness of psychometric effects to inter-individual variability. a**, Relative insensitivity of tested participants to subtle changes in true hazard rate across blocks. Perceived hazard rate estimates in the cue-based (left) and outcome-based (middle) conditions as a function of the true/generative hazard rate (low: more stable blocks, high: more volatile blocks), marked as black lines. Black dots and error bars indicate group-level means ± s.e.m., whereas colored dots indicate participant-level estimates. The perceived hazard rate does not adapt to changes in true hazard, despite the fact that the optimal (accuracy-maximizing) hazard rate (right) tracks changes in true hazard rate. **b**, Left: correlation between perceived hazard rate estimates in more stable (x-axis) and more volatile (y-axis) blocks. Right: correlation between inference noise estimates in more stable and more volatile blocks. Dots and error bars indicate posterior means ± s.d. obtained by model fitting. The thin dotted line shows the identity line. Both parameters correlate strongly between blocks across tested participants. **c**, Psychometric effects for participants with more volatile inference ($N$ = 12). Left: perceived hazard rate estimates. Middle: response repetition curves, showing a clear leftward shift in the outcome-based condition. Lines indicates group-level means ± s.e.m. Right: best-fitting psychometric parameters. The PSE (left) is increased in the outcome-based condition, whereas the sensitivity to evidence (right) is equal across conditions. **d**, Psychometric effects for participants with more stable inference ($N$ = 12). Left: perceived hazard rate estimates. Middle: response repetition curves, showing also a clear leftward shift in the outcome-based condition. Right: best-fitting psychometric parameters. Like the other group, the PSE (left) is increased in the outcome-based condition, whereas the sensitivity to evidence (right) is equal across conditions. Two stars indicate a significant effect at $p < 0.01$, three stars at $p < 0.001$, *n.s.* a non-significant effect.

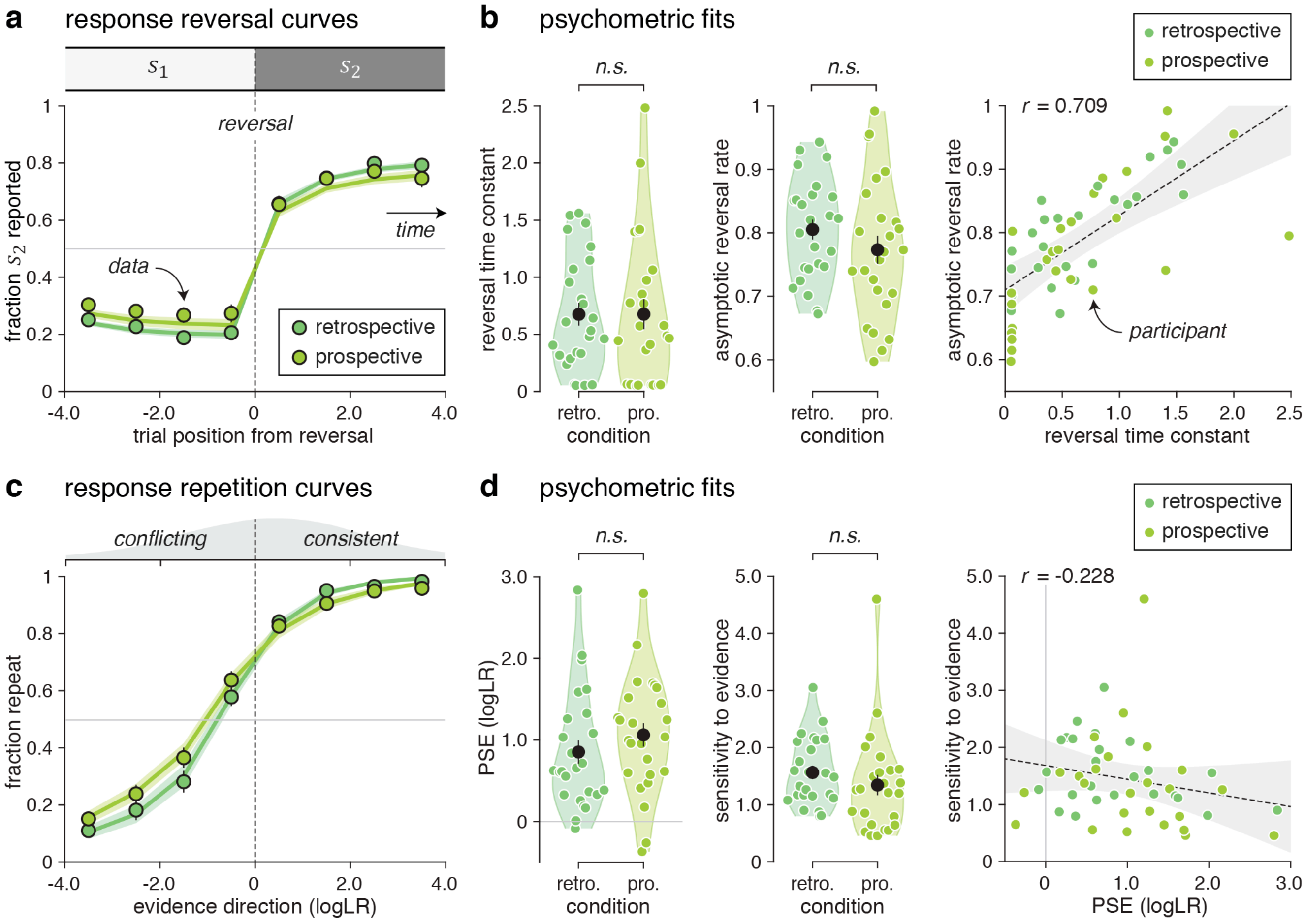


**Supplementary Fig. 5 – Reversal learning behavior during retrospective and prospective cue-based inference.** Results obtained from an additional behavioral dataset ($N$ = 25) in the cue-based condition where participants are instructed to report either the category of the current sequence (retrospective condition) or the next sequence (prospective condition). **a**, Response reversal curves. The thin dotted line indicates the position of the reversal. Dots indicate the observed data (means ± s.e.m.), whereas lines indicate best-fitting saturating exponential functions. **b**, Best-fitting parameters of saturating exponential functions in the retrospective and prospective conditions. Left: the reversal time constant is not slower and the asymptotic reversal rate is not higher in the prospective condition. Black dots and error bars indicate group-level means ± s.e.m., whereas colored dots indicate participant-level estimates. Right: correlation between psychometric parameters. As in the main experiment, the reversal time constant and asymptotic reversal rate correlate positively across tested participants. **c**, Response repetition curves. The thin dotted line indicates perfectly uncertain (null) evidence. Lines indicate best-fitting sigmoid functions. **d**, Best-fitting parameters of sigmoid functions in the retrospective and prospective conditions. Left: the PSE does not increase in the prospective condition. Right: correlation between psychometric parameters. As in the main experiment, the PSE and sensitivity to evidence correlate weakly across tested participants. *n.s.* indicates a non-significant effect.

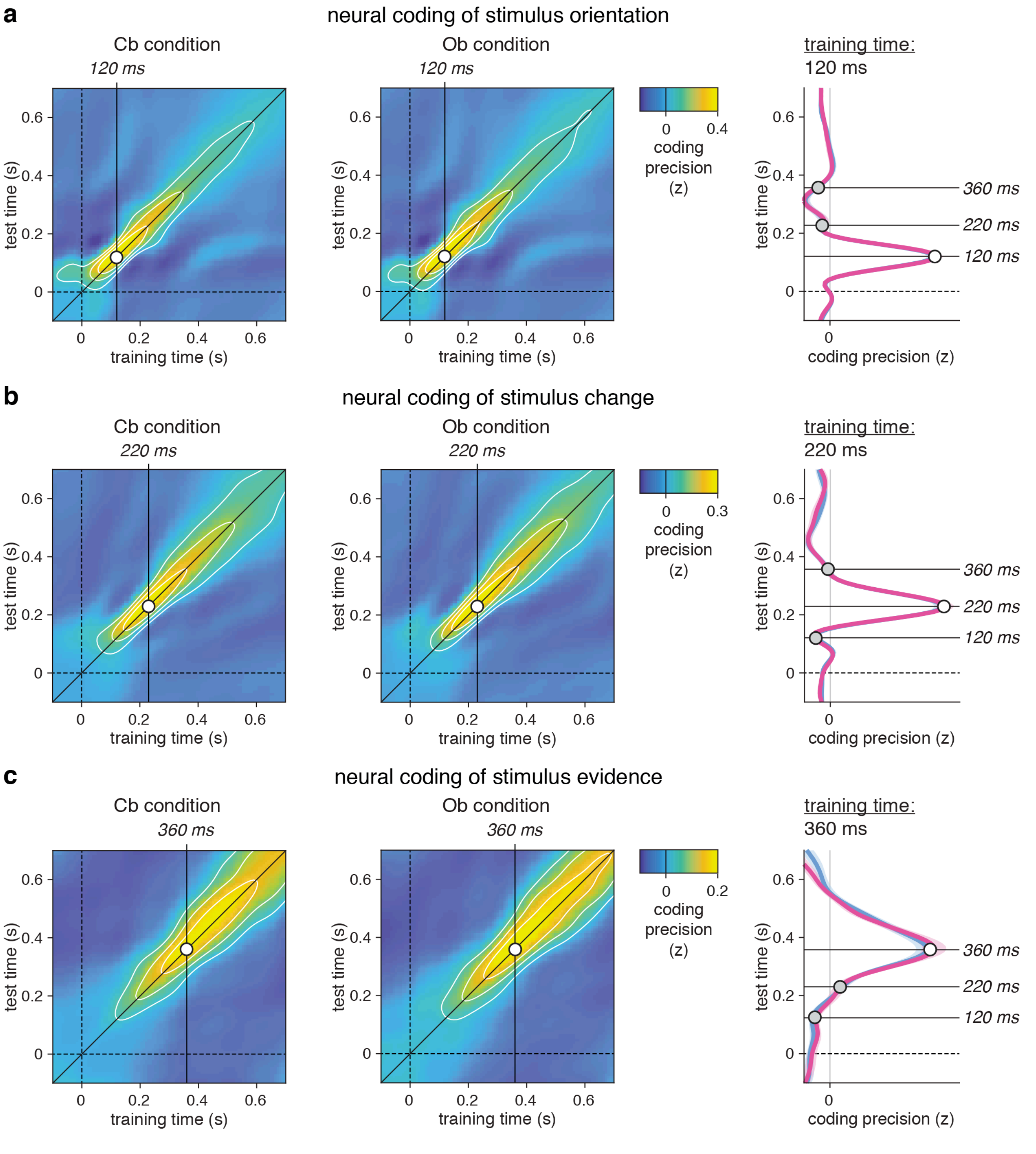


**Supplementary Fig. 6 – Cross-temporal generalization matrices of stimulus characteristics. a**, Cross-temporal generalization of stimulus orientation from whole-brain MEG sensors. Left: training-test matrices obtained in the cue-based (left) and outcome-based (right) conditions. Right: cross-test section trained at the time point associated with maximum coding precision (120 ms). Stimulus orientation is associated with a highly dynamic (sharp diagonal) representation in MEG signals. Lines and shaded error bars indicate group-level means ± s.e.m. **b**, Cross-temporal generalization of stimulus change from whole-brain MEG sensors. Left: training-test matrices in the cue-based and outcome-based conditions. Right: cross-test section trained at the time point associated with maximum coding precision (220 ms). Stimulus change is also associated with a highly dynamic (sharp diagonal) representation in MEG signals. **c**, Cross-temporal generalization of stimulus evidence from whole-brain MEG sensors. Left: training-test matrices in the cue-based and outcome-based conditions. Right: cross-test section trained at the time point associated with maximum coding precision (360 ms). Stimulus evidence is associated with a slightly less dynamic, more stable (wider diagonal) representation in MEG signals.

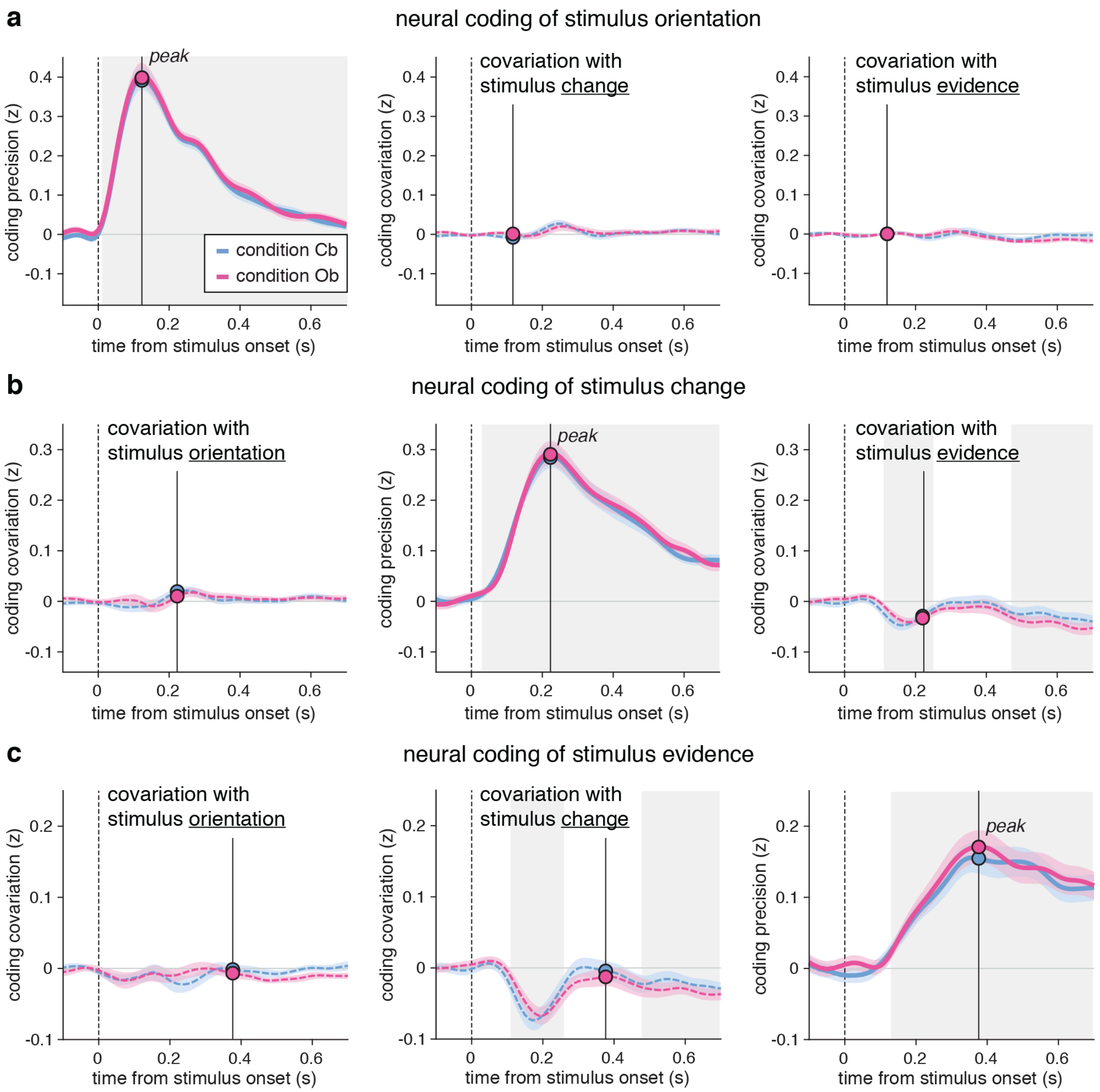


**Supplementary Fig. 7 – Cross-characteristic covariations of neural codes. a**, Cross-characteristic covariations of the neural coding of stimulus orientation from whole-brain MEG sensors. Left: cross-validated correlation between the MEG data projected on the dimension coding for stimulus orientation $\hat{x}_{\text{ori}}$ and the true stimulus orientation. Coding precision peaks at 120 ms following stimulus onset. Middle: correlation between $\hat{x}_{\text{ori}}$ and stimulus change. Right: correlation between $\hat{x}_{\text{ori}}$ and stimulus evidence. The neural code of stimulus orientation evolves through 'null' dimensions for the other two characteristics. **b**, Cross-characteristic covariations of the neural coding of stimulus change from whole-brain MEG sensors. Left: correlation between the MEG data projected on the dimension coding for stimulus change $\hat{x}_{\text{ch}}$ and stimulus orientation. Middle: correlation between $\hat{x}_{\text{ch}}$ and the true stimulus change. Coding precision peaks at 220 ms following stimulus onset. Right: correlation between $\hat{x}_{\text{ch}}$ and stimulus evidence. The neural code of stimulus change evolves through mostly null dimensions for the other two characteristics. **c**, Cross-characteristic covariations of the neural coding of stimulus evidence from whole-brain MEG signals. Left: correlation between the MEG data projected on the dimension coding for stimulus evidence $\hat{x}_{\text{evi}}$ and stimulus orientation. Middle: correlation between $\hat{x}_{\text{evi}}$ and stimulus change. Right: correlation between $\hat{x}_{\text{evi}}$ and the true stimulus evidence. Coding precision peaks at 360 ms following stimulus onset. The neural code of stimulus evidence evolves through mostly null dimensions for the other two characteristics, except around 200 ms following stimulus onset where it covaries negatively with stimulus change. Lines and shaded error bars indicate group-level means ± s.e.m. Shaded gray areas indicate significant effects (cluster-corrected $p < 0.001$).

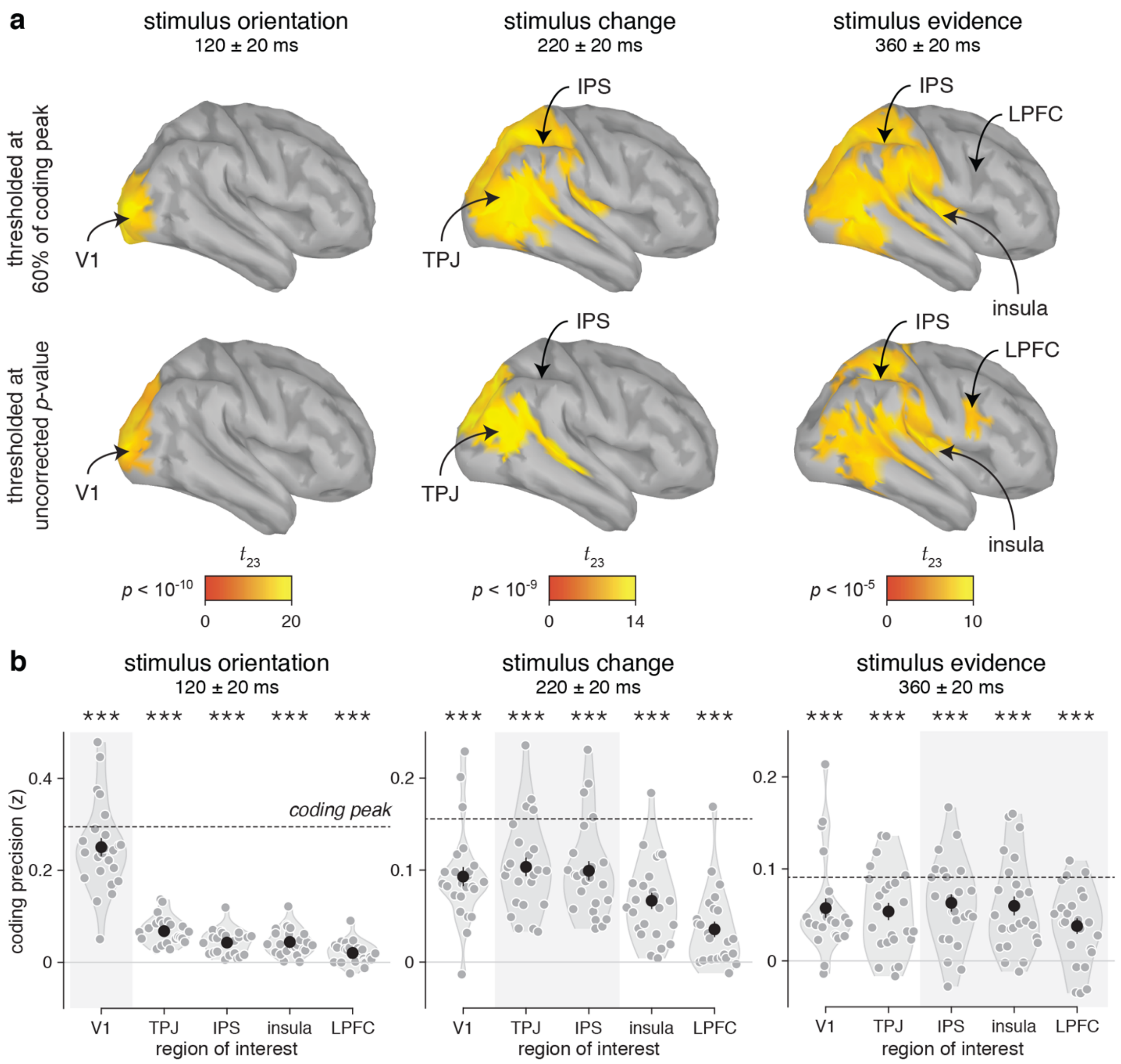


**Supplementary Fig. 8 – Searchlight-based coding of stimulus characteristics across the cortical surface. a**, Searchlight-based coding precision of stimulus characteristics (left column: orientation, middle column: change, right column: evidence) at the latency of their peak, thresholded at 60% of their coding peak (top row) or at an arbitrary, uncorrected *p*-value (bottom row). The neural coding of stimulus orientation peaks in occipital cortex surrounding V1. The neural coding of stimulus change peaks in parietal cortex surrounding the IPS, and in the TPJ. The neural coding of stimulus evidence overlaps broadly with the neural coding of stimulus change, but additionally includes the insula and the LPFC. **b**, Neural coding precision of stimulus characteristics (left: orientation, middle: change, right: evidence) at the latency of their peak across five regions-of-interest. All three stimulus characteristics are coded significantly above chance in all five regions-of-interest. Black dots and error bars indicate group-level means ± s.e.m., whereas colored dots indicate participant-level estimates. Dashed lines indicate the global coding peak of each stimulus characteristic across all cortical sources. Abbreviations: V1 for primary visual cortex, TPJ for temporoparietal junction, IPS for intraparietal sulcus, and LPFC for lateral prefrontal cortex. Three stars indicate a significant effect at $p < 0.001$.

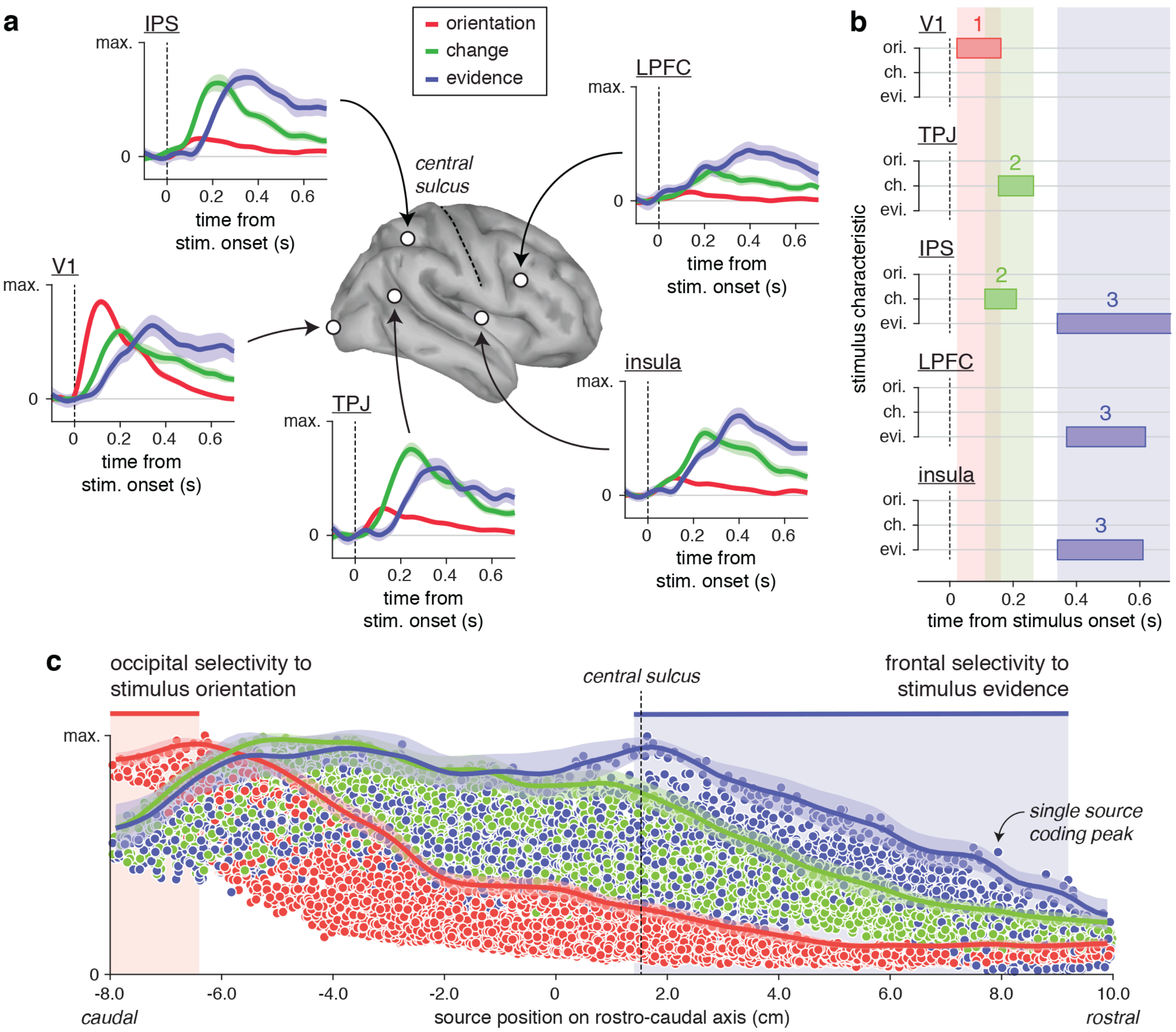


**Supplementary Fig. 9 – Searchlight-based selectivity to stimulus characteristics across the cortical surface. a,** Normalized time courses of neural coding of stimulus orientation (red), stimulus change (green) and stimulus evidence (blue) in five regions-of-interest. The neural coding of each stimulus characteristic is normalized by its global coding across time points and cortical sources to correct for large differences between stimulus characteristics. Selectivity is defined as higher normalized coding of a stimulus characteristic than the other two characteristics. All five regions-of-interest show simultaneous coding of the three stimulus characteristics, but to different extents over time. Lines and shaded error bars indicate jackknifed group-level means ± s.e.m. **b**, Temporal clusters of selectivity to stimulus characteristics in the five regions-of-interest, corrected for multiple comparisons at the cluster level at $p < 0.001$. V1 shows early selectivity to stimulus orientation at 100 ms following stimulus onset. The TPJ shows later selectivity to stimulus change at 200 ms. The IPS shows selectivity to stimulus change at 200 ms, and selectivity to stimulus evidence after 350 ms. The LPFC and insula show selectivity to stimulus evidence after 350 ms. **c**, Rostro-caudal gradient of selectivity to stimulus characteristics. Dots indicate peaks of normalized coding of each stimulus characteristic (red: orientation, green: change, blue: evidence) for each cortical source ($N$ = 5,000) along the rostro-caudal axis, from occipital cortex (left) to frontal cortex (right). Lines and shaded error bars indicate jackknifed means ± s.e.m. of coding peaks for each stimulus characteristic along the rostro-caudal axis, obtained using robust spline smoothing. Coding peaks show a rostro-caudal gradient of selectivity to stimulus characteristics, from occipital selectivity to stimulus orientation (red-shaded area on the left) to frontal selectivity to stimulus evidence (blue-shaded area on the right). The vertical dashed line indicates the position of the central sulcus on the rostro-caudal axis. Abbreviations: V1 for primary visual cortex, TPJ for temporoparietal junction, IPS for intraparietal sulcus, and LPFC for lateral prefrontal cortex.

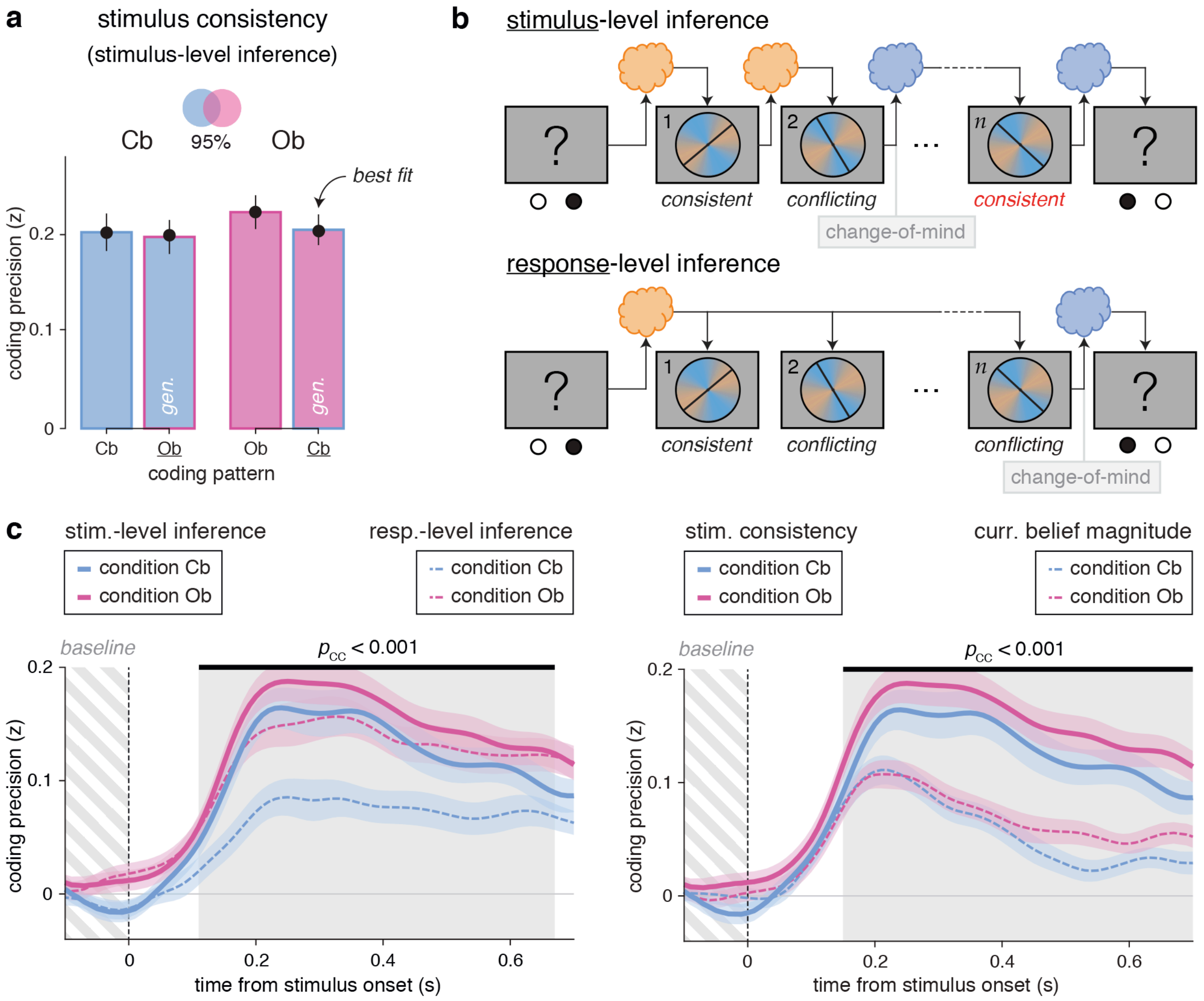


**Supplementary Fig. 10 – Neural evidence for stimulus-level hidden-state inference.** **a**, Estimated coding similarity for stimulus consistency across conditions. Cross-condition generalization indicates near-perfect similarity across conditions (jack-knifed mean: 95%). Bars and error bars indicate group-level means ± s.e.m. Black dots show best-fitting values from the similarity estimation procedure. **b**, Alternative update schemes for hidden-state inference indistinguishable from behavior alone. Top row: stimulus-level inference. This scheme assumes that participants update their belief in the current value of the hidden state after each stimulus, throughout each sequence of stimuli. In this same example, the change-of-mind now occurs midway through the sequence, as soon as a conflicting stimulus (here, stimulus 2) flips the sign of the log-odds belief. Under this stimulus-level update scheme, stimulus consistency is defined as the evidence provided by each stimulus in favor of the current belief accounting for previous stimuli in the same sequence. Bottom row: response-level inference. This scheme assumes that participants update their belief in the current value of the hidden state when probed for a response, following each sequence of stimuli. In this example, a change-of-mind occurs when participants combine their prior belief with the evidence provided by the current sequence. Under this response-level update scheme, stimulus consistency is defined as the evidence provided by each stimulus in favor of the previous response – which reflects the prior belief at the beginning of the current trial. Positive consistency indicates evidence consistent with the previous response (such as stimulus 1), whereas negative consistency indicates evidence conflicting with the previous response (such as stimuli 2 and $n$). As can be seen, the two alternative definitions of stimulus consistency differ for all stimuli presented after a mid-sequence belief switch – such as stimulus $n$ in this example. **c**, Left: neural coding of stimulus consistency assuming stimulus-level inference (solid lines) and response-level inference (dashed lines). Stimulus consistency is coded more precisely in relation to the current belief (stimulus-level inference) than to the prior belief (response-level inference). The shaded area indicates the significant difference in coding precision between the two update schemes. Lines and shaded error bars indicate group-level means ± s.e.m. of coding precision, baseline-corrected across conditions using the last 100 ms preceding stimulus onset (dashed area) to account for possible non-zero correlations across successive stimuli. Right: neural coding of stimulus consistency (solid lines) and the magnitude of the current belief (dashed lines). Stimulus consistency is coded in a stronger and more sustained fashion than the magnitude of the current belief. The shaded area indicates the significant difference in coding precision between the two quantities.

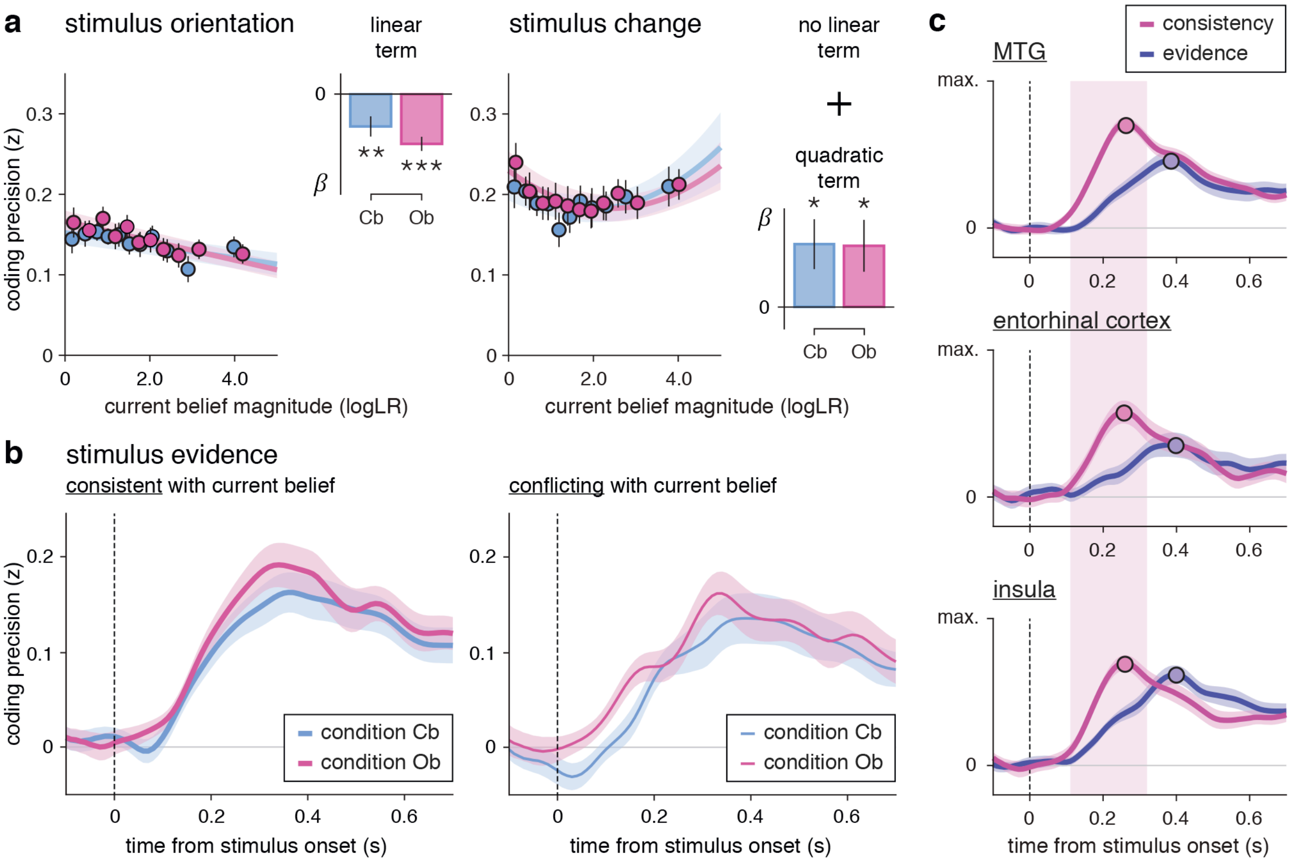


**Supplementary Fig. 11 – Neural dissociation between absolute and relational coding of evidence. a**, Effect of current belief magnitude on the neural coding of stimulus orientation (left) and stimulus change (right), grouped in equally sized bins. Like stimulus evidence, the coding precision of stimulus orientation (left) scales slightly negatively with belief magnitude. The coding precision of stimulus change (right) shows a U-shaped relationship with belief magnitude. Dots and error bars indicate group-level means ± s.e.m. Lines and shaded error bars show least-squares parametric fits. Insets: linear term (stimulus orientation) and quadratic term (stimulus change) estimates (group-level means ± s.e.m) of coding precision against current belief magnitude. **b**, Time course of coding precision of stimulus evidence for stimuli consistent with the current belief (left) and stimuli conflicting with the current belief (right). The neural coding of stimulus evidence does not differ between conditions for consistent nor conflicting stimuli. Lines and error bars indicate group-level means ± s.e.m. **c,** Normalized time courses of neural coding of stimulus consistency (violet) and stimulus evidence (blue) in three regions-of-interest. The MTG, entorhinal cortex and insula show selectivity to stimulus consistency around 200 ms following stimulus onset. Lines and shaded error bars indicate jackknifed group-level means ± s.e.m. Abbreviation: MTG for middle temporal gyrus.

### Computational model specifications

#### Generative model of the task

The generative model of the task is as follows. Each trial $t$ in a sequence of $T$ trials has an associated hidden state $s_t \in \{1,2\}$ that the decision-maker aims to infer. Across successive trials, the hidden state remains unchanged with probability $1-h$ and changes with probability $h$ (hazard rate), that is $p(s_t = s_{t-1}|z_{t-1}) = 1-h$. In each trial $s_t$ the decision-maker gains information about the current $s_t$ by observing the sample sequence $s_{t,1:n_t}$ of $n_t$ oriented patterns, drawn independent and identically distributed (i.i.d.) by dividing a draw from a von Mises distribution with mean $\mu_{s_t}$ (i.e., either $\mu_1$ or $\mu_2$) and concentration $\kappa$ by two. That is, the likelihood of each sample $\theta_{t,i}$ is given by

$$p(\theta_{t,i}|s_t) \equiv \frac{e^{\kappa\cos\left(2(\theta_{t,i}-\mu_{s_t})\right)}}{\pi I_0(\kappa)} \quad (1)$$

where $I_0(\cdot)$ is the zero-order modified Bessel function. Overall, this leads to the following generative model of the task:

$$\begin{aligned} &s_0 \sim \mathcal{B}(p_{s_0}) \\ &s_t|s_{t-1},\tau \sim \mathcal{B}\left(h^{|s_t-s_{t-1}|}(1-h)^{1-|s_t-s_{t-1}|}\right) \\ &\theta_{t,i}|s_t,\kappa,\mu_{1:2} \sim {}^1\!/_2\, \mathcal{VM}(2\,\mu_{s_t},\kappa) \end{aligned} \quad (2)$$

where $\mathcal{B}(\cdot)$ denotes a Bernoulli random variable, $\mathcal{VM}(\cdot,\cdot)$ a von Mises random variable, and $p_{s_0}$ the a-priori belief that $s_0 = 1$.

This model relates to the two experimental conditions as follows. In the cue-based condition, the hidden state $s_t$ identifies the category (A or B) associated with the stimulus sequence $\Theta_t \equiv \{\theta_{t,1:n_t}\}$ observed in each trial. In the outcome-based condition, the hidden state $s_t$ describes the mapping between the response key (left or right) and the category (A or B) of the stimulus sequence it generates. The left response key draws stimulus sequences from category A and the right key from category B if $s_t = 1$, and vice versa if $s_t = 2$. In both conditions, the stimulus sequence $\Theta_t$ at trial $t$ informs about the current hidden state $s_t$.

#### Bayes-optimal decision-making

Let us first derive the Bayes-optimal noise-free decision-making strategy without any bias. Let $g_t \equiv p(s_t = 1|\Theta_{1:t-1})$ denote the prior belief that $s_t = 1$ in the $t^{\text{th}}$ trial after having observed stimulus sequences in trials 1 to $t-1$. Upon observing the stimulus sequence $\theta_{t,1:n_t}$, this belief is updated by Bayes' rule, resulting in

$$\tilde{g}_t \equiv p(s_t = 1|\Theta_{1:t}) \propto g_t \prod_{i=1}^{n_t} p(\theta_{t,i}|s_t = 1) \propto \frac{1}{1+e^{-\left(\kappa\,\mathcal{L}_t + \log\frac{g_t}{1-g_t}\right)}} \quad (3)$$

where all proportionalities are with respect to $s_t$, we have used the above expression for the likelihood, and have defined the sample sequence evidence for $s_t = 1$ by

$$\mathcal{L}_t = \sum_{i=1}^{n_t} \kappa \left( \cos\left(2(\theta_{t,i} - \mu_1)\right) - \cos\left(2(\theta_{t,i} - \mu_2)\right) \right) \tag{4}$$

If we define the prior log-odds belief by $L_t \equiv \log(g_t/(1-g_t))$, then the update simplifies to

$$\tilde{L}_t = L_t + \mathcal{L}_t \tag{5}$$

We find the prior belief $g_{t+1}$ for the next trial $t+1$ from the posterior belief $\tilde{g}_t$ at trial $t$ by the following marginalization:

$$g_{t+1} = \sum_{s_t \in \{1,2\}} p(s_{t+1} = 1 | s_t) p(s_t | \Theta_{1:t}) = h(1 - \tilde{g}_t) + (1-h)\tilde{g}_t \tag{6}$$

In log-odds, this corresponds to

$$L_{t+1} = \mathcal{F}(\tilde{L}_t) \equiv \tilde{L}_t + \log\left(\frac{1-h}{h} + e^{-\tilde{L}_t}\right) - \log\left(\frac{1-h}{h} + e^{+\tilde{L}_t}\right) \tag{7}$$

Bayes-optimal inference would initially use $\tilde{g}_0 = p_{s_0}$, and then alternate between the two above steps to update the belief across successive trials.

Optimal decisions in the cue-based condition correspond to choosing according to $\tilde{g}_t$ in the $t^{\text{th}}$ trial, which is the belief after having observed the corresponding stimulus sequence. $\tilde{g}_t \geq 1/2$ implies that $s_t = 1$ is more likely than $s_t = 2$, and should trigger the action associated with $s_t = 1$. $\tilde{g}_t < 1/2$ implies the opposite, and thus should trigger the action associated with $s_t = 2$. In the outcome-based condition, the relevant belief that should trigger the action in the $t^{\text{th}}$ trial is whether $g_{t+1} \geq 1/2$ or $g_{t+1} < 1/2$. However, by the above update equations we can guarantee that $g_{t+1} \geq 1/2$ as long as $\tilde{g}_t \geq 1/2$ (and $h < 1/2$), such that we can equally cast choices based on the value of $\tilde{g}_t$ – as in the cue-based condition. Thus, we can use the same Bayes-optimal decision-making model to analyze both conditions on equal footing.

#### Bayes-optimal decision-making corrupted by noise and biases

We assume noise arising at three points of the inference process: noise $\sigma_{\text{inf}}$ during inference itself (i.e., the cognitive process of accumulating the evidence provided by stimuli in each trial), noise $\sigma_{\text{sel}}$ during action selection, and transition noise $\sigma_{\text{tr}}$ when computing $\mathcal{F}(\cdot)$ to transition from $\tilde{L}_t$ to $L_{t+1}$. In addition to noise, we assume four biases: 1. an inference leak $\gamma$ while processing the stimulus sequence, 2. a response bias $b$, 3. response lapses occurring with probability $p_{\text{lapse}}$, and 4. an alternative heuristic $\mathcal{G}(\cdot)$ to the Bayes-optimal transition function $\mathcal{F}(\cdot)$. Let us discuss below their exact formulations, in turn.

Starting with the leak $\gamma$, we assume the accumulation of evidence provided by individual stimuli within a sequence to be leaky, which we formulate by changing $\mathcal{L}_t$ to

$$\mathcal{L}_t^{\gamma} = \sum_{i=1}^{n_t} \kappa\, \gamma^{n_t - i} \left(\cos\left(2(\theta_{t,i} - \mu_1)\right) - \cos\left(2(\theta_{t,i} - \mu_2)\right)\right) \tag{8}$$

This corresponds to, at the onset of each stimulus $\theta_{t,i}$, down-weighting (for $\gamma < 1$) or up-weighting (for $\gamma > 1$) of the evidence accumulated so far by $\gamma$. Bayes-optimal evidence accumulation corresponds to $\gamma = 1$.

Inference noise $\sigma_{\text{inf}}$ reflects any noise in the inference process during evidence accumulation. As we have previously shown[1], accumulating $n_t$ stimuli introduces $n_t$ samples of random noise. If each of these noise samples is normally distributed with zero mean and variance $\sigma_{\text{inf}}^2$, then the overall noise added at the end of a stimulus sequence would have variance $n_t\, \sigma_{\text{inf}}^2$. However, due to reweighting the evidence accumulated from earlier stimuli by $\gamma$, earlier samples of noise also get reweighted, yielding noise variance $\sigma_{\text{inf}}^2\, \gamma_t$, where $\gamma_t$ is given by

$$\gamma_t = \sum_{i=1}^{n_t} \gamma^{2(n_t - i)} \tag{9}$$

Overall, this turns the deterministic sample processing sum of Eq. (5) into a stochastic draw:

$$\tilde{L}_t \mid \mathcal{L}_t^{\gamma}, L_t \sim \mathcal{N}\left(L_t + \mathcal{L}_t^{\gamma}, \sigma_{\text{inf}}^2\, \gamma_t\right) \tag{10}$$

Transition noise $\sigma_{\text{tr}}$ has a similar effect on the transition function $\mathcal{F}(\cdot)$ by adding a single sample of zero-mean noise with variance $\sigma_{\text{tr}}^2$, turning Eq. (7) into a stochastic draw:

$$L_{t+1} \mid \tilde{L}_t \sim \mathcal{N}\left(\mathcal{F}(\tilde{L}_t), \sigma_{\text{tr}}^2\right) \tag{11}$$

For some model variants, we replace the Bayes-optimal transition function $\mathcal{F}(\cdot)$ by a two-parameter heuristic $\mathcal{G}(\cdot)$

$$\mathcal{G}(\tilde{L}_t) = \zeta\, \tilde{L}_t + \eta\, \text{sign}(\tilde{L}_t) \tag{12}$$

where $\text{sign}(\cdot) \in \{-1, 0, +1\}$ is the sign function, $\zeta \in [-1, +1]$ parameterizes the multiplicative gain of the transition, and $\eta$ the additive bias of the transition. This heuristic transition function $\mathcal{G}(\cdot)$ acts as a drop-in linear replacement for $\mathcal{F}(\cdot)$, while leaving all other model components unchanged.

Selection noise $\sigma_{\text{sel}}$ does not perturb the inference process itself, and only affects action selection. We model such noise by adding $\varepsilon_t \sim \mathcal{N}(0, \sigma_{\text{sel}}^2)$ to each $\tilde{L}_t$. Furthermore, the decision-maker might feature a response bias, which we model by comparing the noisy log-odds belief to a non-zero threshold value $b$ rather than the Bayes-optimal $b^* = 0$. Also, we assume a possible bias toward repeating the previous action $r_{t-1}$, which we model by a lapse bias probability $p_{\text{lapse}}$ with which the decision-maker blindly repeats the previous action rather than choosing according to the current posterior belief. Overall, this leads to response repetition (i.e., $r_t = r_{t-1}$) with probability $p_{\text{lapse}}$, and otherwise responses determined by

$$r_t = \begin{cases} 1 & \text{if } \tilde{L}_t + \varepsilon_t + b \geq 0 \\ 0 & \text{otherwise} \end{cases} \tag{13}$$

The introduction of transition noise introduces a subtle difference between noisy inference in the cue-based and the outcome-based conditions. Indeed, the transition noise is applied *after* action selection in the cue-based condition, whereas it perturbs the belief *before* action selection in the outcome-based condition. Because these perturbations can flip the sign of the log-odds belief, they can theoretically impact action selection in the outcome-based condition. However, in practice, transition noise $\sigma_{\mathrm{tr}}$ was found to be negligible during Bayesian model selection, and we thus focus on models without transition noise ($\sigma_{\mathrm{tr}} = 0$) in the main text.

#### Fitting the model to observed behavior

For a sequence of $T$ trials, fully specified by the sequence of stimuli $\Theta_t$ in each of the $t = 1, \dots, T$ trials, we observed a corresponding sequence of responses, $r_1, \dots, r_T$. We here describe how we found the model parameters $\phi$ that resulted in the best match between the observed response sequence and the response probabilities predicted by the model, $p(r_{1:T} | \Theta_{1:T}, \phi)$. In its full form, the model has parameters $\phi = \{\sigma_{\mathrm{inf}}, \sigma_{\mathrm{sel}}, \sigma_{\mathrm{tr}}, h, \gamma, b, p_{\mathrm{lapse}}\}$ (with $h$ replaced by $\zeta$ and $\eta$ for the heuristic transition function), where we always assume an unbiased initial belief $L_0 = 0$. We also fitted reduced models that remove certain components by fixing their corresponding parameter values to zero. For example, a model without selection noise would correspond to $\sigma_{\mathrm{sel}} = 0$.

We found best-fitting parameter values by sampling from the Bayesian posterior over parameters using particle Monte Carlo Markov Chain (MCMC) methods[2]. These methods use standard MCMC methods to sample from the parameter posterior, but replace computation of the parameter likelihood $p(r_{1:\mathrm{T}} | \Theta_{1:\mathrm{T}}, \phi)$ – not possible in closed form – with a noisy but unbiased approximation of this likelihood by a particle filter. As MCMC method we used the adaptive mixture Metropolis method[3] that adapts its proposal distribution in an initial burn-in period to achieve favorable acceptance ratios. In the remainder of this section, we describe how we approximated the parameter likelihood with a particle filter.

##### What we want to compute

Combining noisy evidence accumulation and transition, two consecutive $\tilde{L}_t$ can be related by

$$p(\tilde{L}_t \mid \tilde{L}_{t-1}, \phi) = \mathcal{N}(\tilde{L}_t \mid \mathcal{F}(\tilde{L}_{t-1}) + \mathcal{L}_t^{\gamma}, \sigma_\tau^2 + \sigma_{inf}^2\, \gamma_t) \tag{14}$$

where the above is implicitly conditional on $\Theta_t$ through $\mathcal{L}_t^{\gamma}$. These $\tilde{L}_t$ predict observed responses $r_t$ according to the following equation:

$$p(r_t = 1 | \tilde{L}_t, r_{t-1}, \phi) = (1 - p_{\mathrm{lapse}}) \int \mathcal{I}(\tilde{L}_t + \varepsilon_t + b \geq 0)\, p(\varepsilon_t)\, \mathrm{d}\varepsilon_t + p_{\mathrm{lapse}}\, \mathcal{I}(r_{t-1} = 1) \tag{15}$$

where $\mathcal{I}(a)$ is the identifier function that is one if $a$ is true, and zero otherwise.

We would like to track the posterior belief $p(\tilde{L}_{1:t} | r_{1:t}, \phi)$ recursively by

$$p(\tilde{L}_{1:t}|r_{1:t},\phi) = p(\tilde{L}_{1:t-1}|r_{1:t-1},\phi)\frac{p(r_t|\tilde{L}_t,r_{t-1},\phi)p(\tilde{L}_t|\tilde{L}_{t-1},\phi)}{p(r_t|r_{1:t-1},\phi)} \tag{16}$$

$$p(r_{1:t}|\phi) = p(r_{1:t-1}|\phi)p(r_t|r_{1:t-1},\phi) \tag{17}$$

where

$$p(r_t|r_{1:t-1},\phi) = \iint p(r_t|\tilde{L}_t,r_{t-1},\phi)\, p(\tilde{L}_t|\tilde{L}_{t-1},\phi)\, p(\tilde{L}_{t-1}|r_{1:t-1},\phi)\, \mathrm{d}\tilde{L}_t\, \mathrm{d}\tilde{L}_{t-1} \tag{18}$$

Given the above model assumptions, this is unfortunately not possible in closed form. Therefore, we will approximate it by a particle filter.

**Particle filter specification**

The particle filter maintains $K$ particle trajectories $L_{1:t}^1, \ldots, L_{1:t}^K$ that in combination approximate $p(\tilde{L}_{1:t}|r_{1:t},\phi)$. It relies on introducing the importance densities $q(\tilde{L}_1|r_{0:1},\phi)$ for the first trial, and $q(\tilde{L}_t|\tilde{L}_{t-1},r_t,\phi)$ for all trials thereafter, which are used to sample the particles across trials (see next section, **Sampling the importance densities**). Here, $r_0$ is the response provided before any evidence has been presented. The associated importance weights are given by

$$w_1(\tilde{L}_1) = \frac{p(r_1|\tilde{L}_1,r_0,\phi)\, p(\tilde{L}_1|\phi)}{q(\tilde{L}_1|r_{0:1},\phi)} \tag{19}$$

$$w_n(\tilde{L}_{t-1:t}) = \frac{p(r_t|\tilde{L}_t,r_{t-1},\phi)\, p(\tilde{L}_t|\tilde{L}_{t-1},\phi)}{q(\tilde{L}_t|\tilde{L}_{t-1},r_{t-1:t},\phi)} \tag{20}$$

The particle filter operates as follows. In the first trial, it samples $L_1^k \sim q(\tilde{L}_1|r_1,\phi)$, computes the associated weights $w_1(L_1^k)$, and sets $W_1^k \propto w_1(L_1^k)$ such that $\sum_k W_1^k = 1$. It then resamples the $L_1^k$'s according to the weights $W_1^k$ to obtain $K$ equally-weighted particles $\bar{L}_1^k$. For all further trials, $t \geq 2$, the particle filter first samples $L_t^k \sim q(\tilde{L}_t|\bar{L}_{t-1}^k,r_{t-1:t},\phi)$ and sets $L_{1:t}^k \leftarrow (\bar{L}_{1:t-1}^k L_t^k)$, where $\bar{L}_{1:t-1}^k$ is the trajectory from $1$ to $t-1$ associated with the particle $\bar{L}_1^{t-1}$. Based on these samples, it computes the weights $w_t(L_{t-1:t}^k)$ and sets $W_t^k \propto w_t(L_{t-1:t}^k)$ such that $\sum_k W_t^k = 1$. It the resamples the $L_t^k$ according to the weights $W_t^k$ to obtain $K$ equally-weighted particles $\bar{L}_t^k$.

After $t$ trials, the densities $p(\tilde{L}_{1:t}|r_{1:t},\phi)$ and $p(r_t|r_{1:t-1},\phi)$ are approximated by

$$\hat{p}(d\tilde{L}_{1:t}|r_{1:t},\phi) = \sum_k W_t^k \delta_{L_{1:t}^k}(d\tilde{L}_{1:t}) \tag{21}$$

$$\hat{p}(r_t|r_{1:t-1},\phi) = \frac{1}{K}\sum_k w_t(L_{t-1:t}^k) \tag{22}$$

We use the latter to estimate the marginal likelihood $p(r_{1:T}|\phi)$ by

$$\hat{p}(r_{1:T}|\phi) = \prod_{t=1}^{T} \hat{p}(r_t|r_{1:t-1}, \phi) \tag{23}$$

This estimate is unbiased, and thus suitable for use in MCMC methods[2].

**Sampling the importance densities**

What remains is to define the importance densities and to describe how to sample from them. The optimal choice for these densities is given by

$$q(\tilde{L}_1|r_{0:1}, \phi) = p(\tilde{L}_1|r_{0:1}, \phi) \propto p(r_1|\tilde{L}_1, r_0, \phi)p(\tilde{L}_1|\phi) \tag{24}$$

$$q(\tilde{L}_t|\tilde{L}_{t-1}, r_{t-1:t}, \phi) = p(\tilde{L}_t|\tilde{L}_{t-1}, r_{t-1:t}, \phi) \propto p(r_t|\tilde{L}_t, r_{t-1}, \phi)p(\tilde{L}_t|\tilde{L}_{t-1}, \phi) \tag{25}$$

Let us now consider the first and the remaining trials separately.

*Particle sampling and weights for the first trial of each block*

To model response lapses, we draw $z_1 \sim \mathcal{B}(p_{\text{lapse}})$, and declare the trial a lapse trial if $z_1 = 1$. For lapse trials, the choice $r_1$ is not informative about $\tilde{L}_1$, such that

$$\tilde{L}_1|(z_1 = 1) \sim \mathcal{N}(\kappa\, \mathcal{L}_1^{\gamma}, \sigma_{\text{inf}}^2 \gamma_1), \tag{26}$$

where we have assumed $L_0 = 0$, as discussed above. For non-lapse trials, where $z_1 = 0$ (implicit in the below notation), $r_1$ stochastically constraints the value of $\tilde{L}_1$ through the likelihood $p(r_1|\tilde{L}_1, \phi)$. To perform tractable sampling from the posterior $p(\tilde{L}_1|r_1, \phi)$, we split the link between $\tilde{L}_1$ and $r_1$ by introducing the auxiliary variable $d_1 = \tilde{L}_1 + \varepsilon_1 + b$, such that

$$p(\tilde{L}_1|r_1, \phi) = \int p(\tilde{L}_1|d_1, \phi)\, p(d_1|r_1, \phi)\, \mathrm{d}d_1. \tag{27}$$

This shows that we can sample $\tilde{L}_1|r_1$ by first sampling $d_1|r_1$, and then $\tilde{L}_1|d_1$. For our model, $p(d_1|r_1, \phi)$ turns out to be given by

$$p(d_1|r_1, \phi) \propto p(r_1|d_1)\, p(d_1|\phi) = \mathcal{N}\left(d_1 \mid \mathcal{L}_1^{\gamma} + b, \sigma_{\text{inf}}^2 \gamma_1 + \sigma_{\text{sel}}^2\right) \begin{cases} \mathcal{I}(d_1 \geq 0) & \text{if } r_1 = 1 \\ \mathcal{I}(d_1 < 0) & \text{otherwise} \end{cases} \tag{28}$$

where we found $p(d_1|\phi)$ by marginalizing $p(d_1|\tilde{L}_1, \varepsilon_1, \phi)\, p(\tilde{L}_1|\phi)\, p(\varepsilon_1|\phi)$ over $\tilde{L}_1$ and $\varepsilon_1$. The above is a truncated normal distribution, for which efficient sampling methods exist.

Given $d_1$, $p(\tilde{L}_1|d_1, \phi)$ becomes normally distributed, and is given by

$$\begin{aligned} p(\tilde{L}_1|d_1, \phi) &\propto p(d_1|\tilde{L}_1, \phi)\, p(\tilde{L}_1|\phi) \\ &\propto \mathcal{N}\left(\tilde{L}_1 \middle| \frac{\sigma_{\text{inf}}^2 \gamma_1}{\sigma_{\text{sel}}^2 + \sigma_{\text{inf}}^2 \gamma_1}(d_1 - b) + \frac{\sigma_{\text{sel}}^2}{\sigma_{\text{sel}}^2 + \sigma_{\text{inf}}^2 \gamma_1} \mathcal{L}_1^{\gamma}, \frac{\sigma_{\text{sel}}^2\, \sigma_{\text{inf}}^2 \gamma_1}{\sigma_{\text{sel}}^2 + \sigma_{\text{inf}}^2 \gamma_1}\right) \end{aligned} \tag{29}$$

which is again easy to sample from.

Given the above importance densities, the particle weights for the first trial are given by $w_1(\tilde{L}_1) = p(r_1|r_0, \phi)$, and are thus independent of the sampled values of $\tilde{L}_1$. Using the $\tilde{L}_1 \to d_1 \to r_1$ split, as before, we find these weights to be given by

$$w_1(\tilde{L}_1) = p(r_1|r_0, \phi) = \sum_{z_1} p(z_1) \int p(r_1|d_1, z_1)\, p(d_1|\phi)\, \mathrm{d}d_1$$

$$= p_{\text{lapse}}\, \mathcal{I}(r_1 = r_0) + (1 - p_{\text{lapse}}) \begin{cases} \Phi\left(\dfrac{\kappa\, \mathcal{L}_1^{\gamma} + b}{\sqrt{\sigma_{\text{sel}}^2 + \sigma_{\text{inf}}^2\, \gamma_1}}\right) & \text{if } r_1 = 1 \\ 1 - \Phi\left(\dfrac{\kappa\, \mathcal{L}_1^{\gamma} + b}{\sqrt{\sigma_{\text{sel}}^2 + \sigma_{\text{inf}}^2\, \gamma_1}}\right) & \text{otherwise} \end{cases} \quad (30)$$

This completes specifying all the components for particle sampling and computing the particle weights for the first trial.

*Particle sampling and weights for all remaining trials*

For all remaining trials of each block ($t \geq 2$), we again draw response lapse trials according to $z_t \sim \mathcal{B}(p_{\text{lapse}})$. If $z_t = 1$, then $r_t$ is uninformative about $\tilde{L}_t$, such that we draw log-odds beliefs using a non-truncated distribution:

$$\tilde{L}_t \mid \tilde{L}_{t-1}, (z_t = 1) \sim \mathcal{N}\left(\mathcal{F}(\tilde{L}_{t-1}) + \mathcal{L}_t^{\gamma}, \sigma_{\text{tr}}^2 + \sigma_{\text{inf}}^2\, \gamma_t\right) \quad (31)$$

For non-lapse trials, when $z_t = 0$ (implicitly conditioned on below), we again introduce the auxiliary variable $d_t = \tilde{L}_t + \varepsilon_t + b$ to draw from $\tilde{L}_t|\tilde{L}_{t-1}, r_t$ using

$$p(\tilde{L}_t|\tilde{L}_{t-1}, r_t, \phi) = \int p(\tilde{L}_t|\tilde{L}_{t-1}, d_t, \phi)\, p(d_t|\tilde{L}_{t-1}, r_t, \theta)\, \mathrm{d}d_t \quad (32)$$

which allows us to first sample $d_t|\tilde{L}_{t-1}, r_t$, and then $\tilde{L}_t|\tilde{L}_{t-1}, d_t$. To find $p(d_t|\tilde{L}_{t-1}, r_t, \phi)$, we first note that the $p(d_t|\tilde{L}_{t-1}, \phi)$ is by marginalization of $p(d_t|\tilde{L}_t, \varepsilon_t, \phi)\, p(\tilde{L}_t|\tilde{L}_{t-1}, \phi)\, p(\varepsilon_t|\phi)$ given by

$$p(d_t|\tilde{L}_{t-1}, \phi) = \mathcal{N}\left(d_t|\mathcal{F}(\tilde{L}_{t-1}) + \mathcal{L}_t^{\gamma} + b, \sigma_{\text{tr}}^2 + \sigma_{\text{inf}}^2\, \gamma_n + \sigma_{\text{sel}}^2\right) \quad (33)$$

Using this, we find $p(d_t|\tilde{L}_{t-1}, r_t, \phi)$ to be given by

$$\begin{aligned} p(d_t|\tilde{L}_{t-1}, r_t, \phi) &\propto p(r_t|d_t)\, p(d_t|\tilde{L}_{t-1}, \phi) \\ &= \mathcal{N}\left(d_n|\mathcal{F}(\tilde{L}_{t-1}) + \mathcal{L}_t^{\gamma} + b, \sigma_{\text{tr}}^2 + \sigma_{\text{inf}}^2\, \gamma_n + \sigma_{\text{sel}}^2\right) \begin{cases} \mathcal{I}(d_t \geq 0) & \text{if } r_t = 1 \\ \mathcal{I}(d_t < 0) & \text{otherwise} \end{cases} \end{aligned} \quad (34)$$

which is again a truncated normal distribution that can be efficiently sampled from.

Given $d_t$, $p(\tilde{L}_t|\tilde{L}_{t-1}, d_t, \phi)$ becomes normally distributed, and is given by

$$\begin{aligned} p(\tilde{L}_t|\tilde{L}_{t-1}, d_t, \phi) &\propto p(d_t|\tilde{L}_t, \phi)\, p(\tilde{L}_t|\tilde{L}_{t-1}, \phi) \\ &= \mathrm{N}\left(\tilde{L}_t \middle| \frac{(\sigma_{\tau}^2 + \sigma_{\text{inf}}^2\, \gamma_t)(d_t - b) + \sigma_{\text{sel}}^2(\mathcal{F}(\tilde{L}_{t-1}) + \mathcal{L}_t^{\gamma})}{\sigma_{\text{sel}}^2 + \sigma_{\text{tr}}^2 + \sigma_{\text{inf}}^2\, \gamma_t}, \frac{\sigma_{\text{sel}}^2(\sigma_{\text{tr}}^2 + \sigma_{\text{inf}}^2\, \gamma_t)}{\sigma_{\text{sel}}^2 + \sigma_{\text{tr}}^2 + \sigma_{\text{inf}}^2\, \gamma_t}\right) \end{aligned} \quad (35)$$

With the above importance densities, the particle weights for trial $t$ are given by $w_t(\tilde{L}_{t-1:t}) = p(r_t|\tilde{L}_{t-1}, r_{t-1}, \phi)$, which evaluates to

$$w_t(\tilde{L}_{t-1:t}) = p(r_t|\tilde{L}_{t-1}, r_{t-1}, \phi) = \sum_{z_t} p(z_t) \int p(r_t|d_t, z_t)\, p(d_t|\tilde{L}_{t-1}, r_{t-1}, \phi)\, \mathrm{d}d_t$$

$$= p_{\text{lapse}}\, \mathcal{I}(r_t = r_{t-1}) + (1 - p_{\text{lapse}}) \begin{cases} \Phi\left(\dfrac{\mathcal{F}(\tilde{L}_{t-1}) + \mathcal{L}_t^{\gamma} + b}{\sqrt{\sigma_{\text{sel}}^2 + \sigma_{\text{tr}}^2 + \sigma_{\text{inf}}^2 \gamma_t}}\right) & \text{if } r_t = 1 \\ 1 - \Phi\left(\dfrac{\mathcal{F}(\tilde{L}_{t-1}) + \mathcal{L}_t^{\gamma} + b}{\sqrt{\sigma_{\text{sel}}^2 + \sigma_{\text{tr}}^2 + \sigma_{\text{inf}}^2 \gamma_t}}\right) & \text{otherwise} \end{cases} \tag{36}$$

This completes specifying all the components for particle sampling and computing the particle weights for all remaining trials after the first of each block.
